# A genome quality control program for the haploid stage of spermatogenesis

**DOI:** 10.64898/2026.07.27.740997

**Authors:** Maiko Kitaoka, Yukiko M. Yamashita

## Abstract

Genome integrity is fundamental to cellular and organismal health, with the germline genome bearing the most profound impact. During spermatogenesis, post-meiotic germ cells, called spermatids, undergo a series of morphological changes over an extended period (a week in fruit flies, two weeks in humans) to yield mature sperm. Having undergone consecutive meiotic divisions, these spermatids lack homologous chromosomes or sister chromatids, critical templates for accurate DNA repair. How genome integrity of spermatids is ensured during this vulnerable yet prolonged period remains poorly understood. In this study, we show that the process of spermatogenesis is specifically compromised to eliminate damaged nuclei upon γ-irradiation. We further find a non-canonical role of histone variant H2Av in the process of spermatid elimination. Our findings demonstrate the presence of a haploid-specific quality control program that ensures the integrity of the sperm genome transmitted to the next generation.

## INTRODUCTION

Gametes are the sole cell type that can transmit genetic information across generations; thus, their genome integrity is of the utmost importance. Paradoxically, the very process of male gametogenesis presents unique challenges to genome integrity (reviewed in (Kitaoka and Yamashita, 2024)). Across many species, the post-meiotic stages of spermatogenesis, called spermiogenesis, are extremely long-lasting to accommodate major morphological changes to produce mature sperm cells with highly compacted DNA and long flagellar tails (Fabian and Brill, 2012; Lindsley and Tokuyasu, 1980; Heller and Clermont, 1963). Critically, this lengthy period of spermiogenesis coincides with the most vulnerable period for the genome: After having undergone meiotic divisions, spermatids do not have homologous chromosomes or sister chromatids, critical templates for faithful DNA repair (i.e., homologous recombination repair). Furthermore, the spermiogenesis program is largely transcriptionally silent and decouples transcription and translation such that cellular responses to insults are likely limited (White-Cooper, 2010; Barreau et al., 2008). How developing post-meiotic spermatids respond to DNA damage, and how nuclei with damaged genomes may be prevented from being passed down to the next generation, remain poorly understood.

*Drosophila* has served as a leading model to study spermatogenesis owing to the well-defined anatomy of their testes: Developing germ cells are spatiotemporally organized in the testis from the germline stem cells through mitotic, meiotic, and post-meiotic phases, and each stage of sperm development can be identified at single-cell resolution (Fig. 1A). Cytokinesis is incomplete during 4 spermatogonial mitotic divisions and 2 meiotic divisions, producing a germ cell cyst in which all 64 spermatids originating from a single stem cell division are connected to each other. After meiosis, the round nuclei of 64 spermatids undergo a striking morphological transformation synchronously in a cyst, yielding 10 µm needle-shaped nuclei over the course of 70-80 hours (Chandley and Bateman, 1962; Lindsley and Tokuyasu, 1980; Fabian and Brill, 2012; Tokuyasu, 1974). Nuclear compaction and elongation accompany cellular morphological changes as the sperm axonemes grow to an ultimate length of 2 mm (Fig. 1B) (Tokuyasu, 1975). This well-defined progression of spermatogenesis in *Drosophila* allows us to investigate how spermatids may respond to DNA damage post-meiosis to ensure the quality of the genome that participates in fertilization.

**Figure 1.**
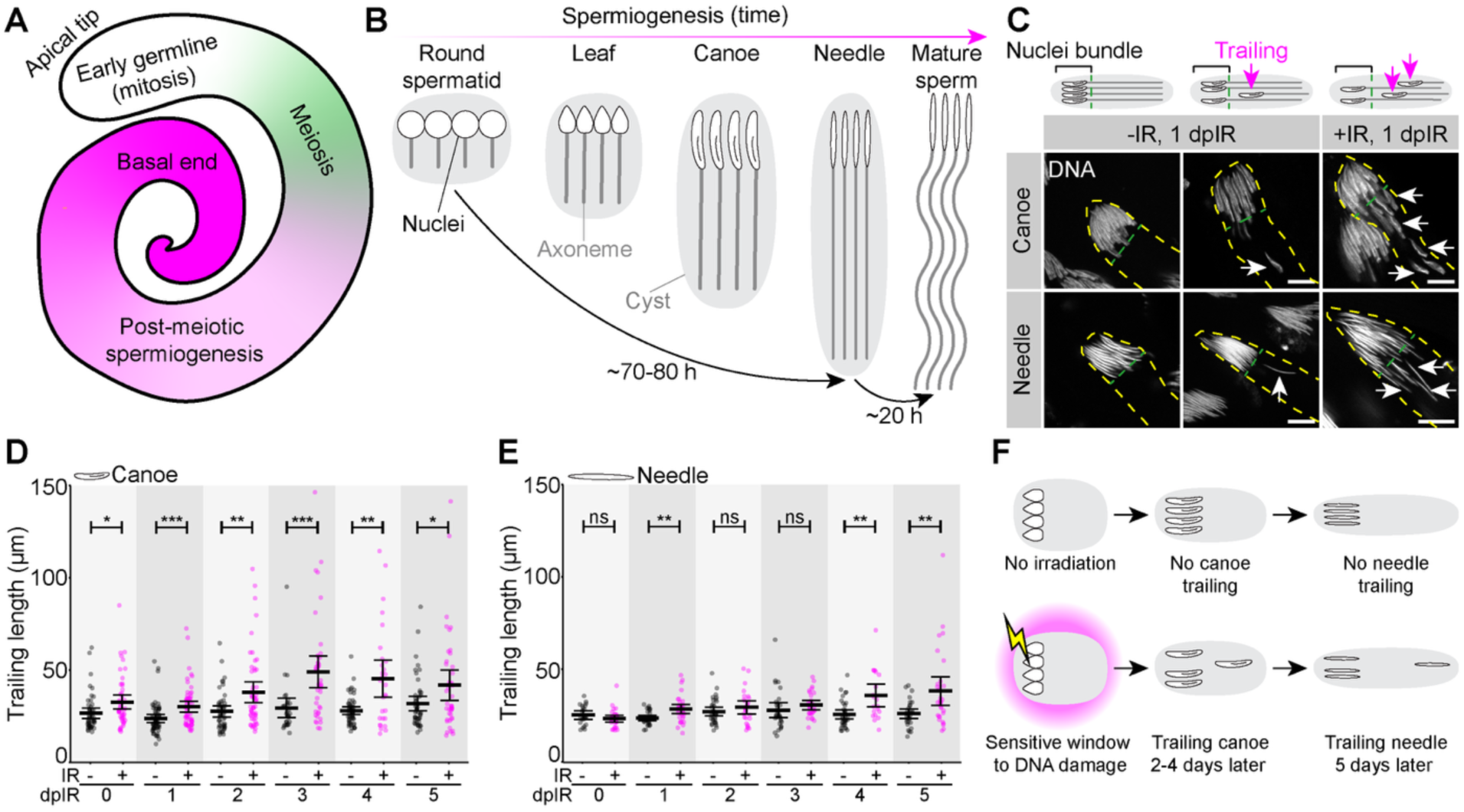
Spermatid nuclei trail after irradiation. **A.** Schematic cartoon of the *Drosophila* testis, highlighting mitotic (white), meiotic (green), and post-meiotic regions (magenta). **B.** Schematic of *Drosophila* post-meiotic spermiogenesis, in particular showing the nuclear morphology staging. **C.** Schematic and representative images of spermatid nuclei stained with DAPI (white) without (left and middle panels) and with irradiation (right panels) at 1 day post irradiation (dpIR). Green dashed lines indicate where the nuclear bundle ends, and arrows indicate trailing nuclei that are separate from the main bundle at the proximal end of the cyst. Yellow dashed lines outline the cyst in each image, scale = 10 µm. **D, E.** Quantification of trailing length in testes without (gray) or with irradiation (magenta) at various timepoints after irradiation for canoe (D) and needle (E) spermatids. The means with 95% CIs are plotted. **F.** Schematic demonstrating the time window when spermatid trailing can be induced.

In this work, we use irradiation-induced DNA damage as a model to investigate how haploid gametes may respond to DNA damage to maintain genome quality for the next generation in the face of the unique challenge of lacking repair templates. We describe a cell biological program that eliminates damaged spermatids during the process of spermiogenesis: We found that, upon γ-irradiation, a subset of spermatids within a cyst were displaced, triggering their elimination and precluding them from participating in fertilization. Surprisingly, this DNA damage-induced elimination of spermatids depends on a non-canonical function of the histone variant H2Av (H2Ax in mammals), which is widely used to detect and signal DNA double-strand breaks (Rogakou et al., 1998; Madigan et al., 2002). Taken together, the present study reveals the presence of a DNA damage response uniquely tailored to eliminate damaged haploid genomes, thereby ensuring the quality of the gamete genome for the next generation.

## RESULTS

### DNA damage by γ-irradiation induces spermatid nuclei trailing

To test how developing spermatids might respond to DNA damage, if at all, we challenged testes with irradiation to induce double-stranded breaks, and examined the response of haploid, post-meiotic spermatids. We focused solely on post-meiotic stages, where cells cannot rely on the canonical DNA damage response, including faithful homologous recombination repair (Edwards and Sirlin, 1958; Lu and Yamashita, 2017; Gunes et al., 2015; Graham et al., 2024). Time points were chosen such that the spermatids to be analyzed were already haploid at the time of irradiation, based on the known time frame of spermatid development (Chandley and Bateman, 1962; Lindsley and Tokuyasu, 1980). The irradiation dose used in this study did not lead to an accumulation of any specific spermatid stage, suggesting that no developmental arrest (akin to cell cycle checkpoints) was induced (Fig. S1A-D). Additionally, the irradiation dose used did not lead to major disorganization of the cysts or testis anatomy (Fig. S1E-F). After examining multiple doses (25 rad to 1000 rad), we chose 500 rad (0.5 Gy) for the rest of the experiments, as this dose was the lowest while reproducibly inducing the stereotypical responses as described below.

Throughout spermatid development, 64 post-meiotic haploid cells differentiate together in a cyst as a bundle, where all haploid cells orient their nuclei toward the basal end of the testis, and their axonemal tails to the opposing apical tip. Nuclear morphology transforms from ‘round’ spermatids immediately after meiosis to ‘leaf’, ‘canoe’, and finally ‘needle’ spermatids (Fig. 1A, B). During the canoe stage, spermatid chromatin transitions from the canonical histone-based packaging to protamine-based packaging, called the histone-to-protamine transition (Rathke et al., 2014, 2007). Prior to this transition, chromatin in the round and leaf stage spermatids still retains histones. Although 64 sperm nuclei are typically closely bundled during differentiation, we observed that many nuclei became separated from the bundle following irradiation (Fig. 1C, D). Here, we define the spermatid ‘bundle’ as a cluster of nuclei that are tightly grouped together and juxtaposed to the most proximal side of the cyst, as observed in the wild type (Fig. 1C), whereas we term the nuclei separated from the bundle as ‘trailing’ spermatid nuclei. Trailing nuclei were observed only occasionally in non-irradiated testes, but increased dramatically upon irradiation (Fig. 1C). As a proxy for the extent of trailing, we measured the distance from the tip of the spermatid bundle to the last visible, identifiable trailing nucleus in the same cyst, as ‘trailing length’. We examined the dynamics of trailing across time points up to 5 days post-irradiation to determine how spermatids respond to irradiation. Interestingly, nuclear trailing seemed to occur in a wave, where canoe spermatid trailing peaked 2-4 days after irradiation, followed by needle spermatid trailing at 5 days after irradiation (Fig. 1D, E). This suggests that there is a sensitive time window before the canoe stage where irradiation can induce spermatid trailing as they progress to canoe and needle stages (Fig. 1F). Based on the known developmental timing (Chandley and Bateman, 1962; Lindsley and Tokuyasu, 1980), this sensitive time window is most likely around the beginning of the haploid stages post-meiosis.

It was sometimes difficult to identify all of the trailing nuclei that belong to a specific cyst as the spermatid tail elongates (>200 µm), and some of the trailing spermatid nuclei that were much farther away from the bundle are likely missed. Thus, the ‘trailing length’ (Fig. 1D, E) is likely an underestimation. As a complementary measure, we counted how many nuclei remained in the bundle after irradiation-induced trailing. To do so, we counted the number of basal bodies instead of nuclei themselves (Methods). The basal body is a centriole-derived structure that plays an essential role in connecting the spermatid nucleus to the flagellar tail (Khire et al., 2016; Buglak et al., 2024), and each nucleus is attached to a single basal body. We found that basal bodies can be accurately counted, whereas the nuclei are difficult to count because they cannot be spatially resolved (Fig. 2A, B). By counting the number of basal bodies within a bundle, we found that the number of nuclei decreases considerably upon irradiation (Fig. 2C), revealing that many nuclei are indeed lost to trailing. Taken together, our results show that irradiation induces trailing of spermatid nuclei and loss of nuclei from the bundle.

**Figure 2.**
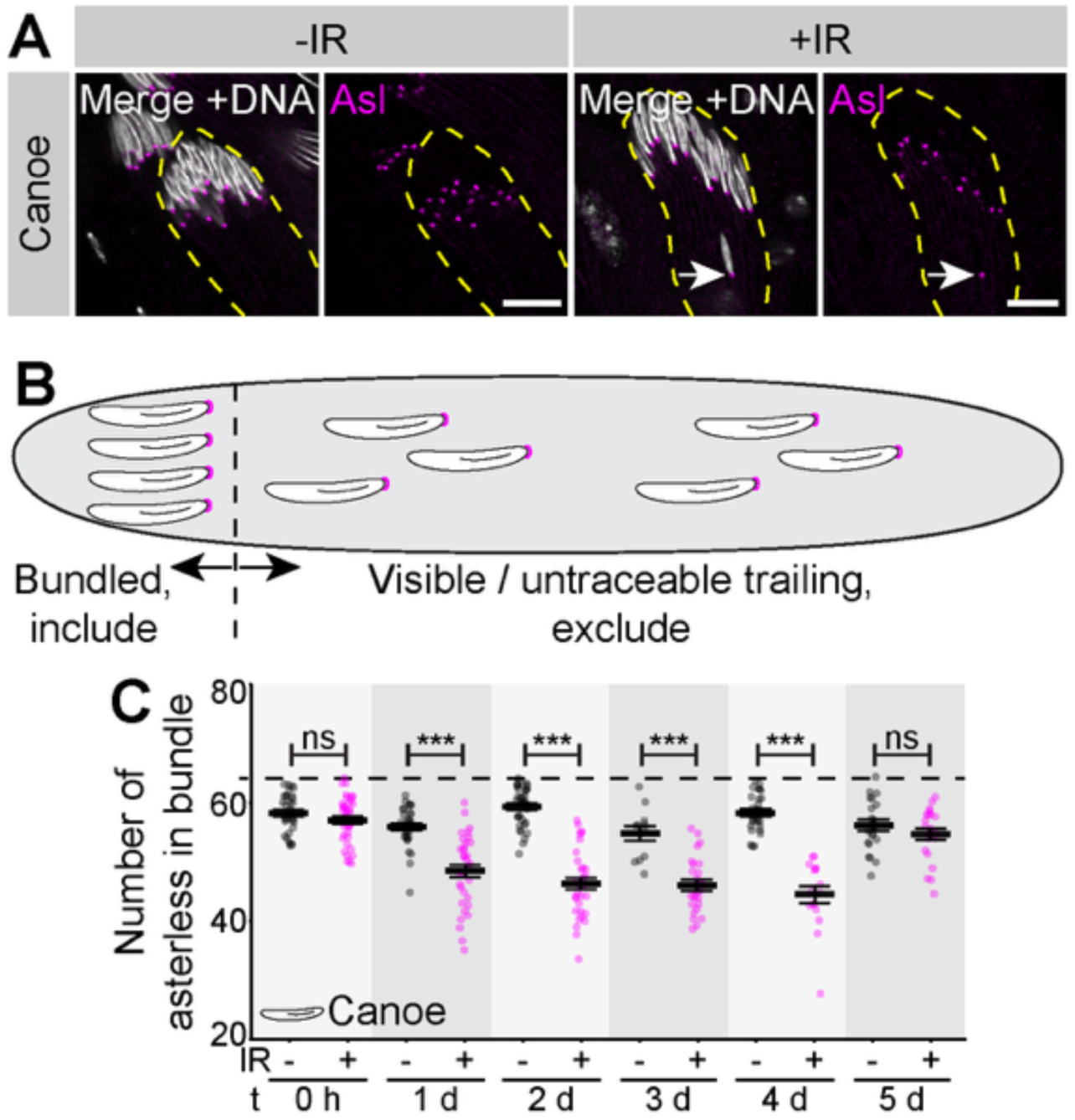
Irradiation induces spermatid loss. **A-B.** Representative images (A) and schematic (B) of canoe spermatids stained with DAPI (white) and asterless (Asl, magenta) to mark the basal body and allow for quantification of spermatids in a bundle. Yellow dashed lines outline the cyst in each image, scale = 10 µm. **C.** Quantification of asterless per bundle in testes without (gray) or with irradiation (magenta) at various time points after irradiation. Dotted line marks 64, the theoretical maximum number of nuclei per cyst. The means with 95% CIs are plotted.

### Trailing spermatid nuclei fail individualization

The trailing nuclei, which fall behind the main bundle, have been assumed to be non-functional, based on the observations that a variety of male sterile mutants often exhibit severe trailing, where essentially all nuclei scatter throughout the length of the cyst (Kimura, 2013; Nerusheva et al., 2009; Napoletano et al., 2017; Couderc et al., 2017; Dorogova et al., 2008; Chen et al., 2021; Herbette et al., 2021; Kracklauer et al., 2010; Yuan et al., 2019). By extension, spermatid nuclei that occasionally trail in wild-type flies (Fig. 1C) were also speculated to be defective. While there are examples of mutants (e.g., axonemal dynein mutants (Fatima, 2011)), where trailing is mechanistically connected to sterility, it remained unclear whether trailing unequivocally leads to dysfunctional sperm.

We found that the trailing nuclei fail during individualization, precluding them from becoming functional sperm. Individualization is a conserved process of spermiogenesis, in which germ cells that have developed as interconnected cells within a cyst become separated from one another, releasing individual sperm cells (Steinhauer, 2015). The trailing spermatids appeared normal until the very late stage of spermiogenesis, irrespective of whether trailing was induced by irradiation or observed sporadically without irradiation. Trailing spermatids progressed normally through nuclear DNA compaction during the histone-to-protamine transition, and through the formation of the individualization complex (IC), an actin-based structure essential for individualization (Fig. 3A-C). The IC is known to remove excess cytoplasm along the entire length of the cyst to separate individual sperm cells from each other (Steinhauer, 2015; Tokuyasu, 1972; Fabrizio et al., 1998; Noguchi and Miller, 2003). However, when they reached the stage of caspase activation, which is known to be required for the individualization process (Arama et al., 2003; Huh et al., 2004; Arama et al., 2006), the ICs of trailing nuclei were lost, while the ICs associated with bundled nuclei were not (Fig. 3D, E). These results suggest that trailing nuclei and their associated IC are not protected from degradation upon caspase activation, unlike the nuclei in the bundle (Arama et al., 2003). In the absence of IC, it is impossible for trailing nuclei to become properly individualized, thus they cannot become functional sperm. Trailing nuclei also became positive for anti-double-stranded DNA (dsDNA) antibody staining at this stage, although normal nuclei at this needle stage are negative for dsDNA due to extreme DNA compaction (Herbette et al., 2021) (Fig. 3E). These results imply that DNA of trailing nuclei is likely cleaved by caspase and decondensed. Our results reveal, for the first time, how trailing spermatid nuclei are eliminated and fail to be passed to the next generation, due to defective individualization.

**Figure 3.**
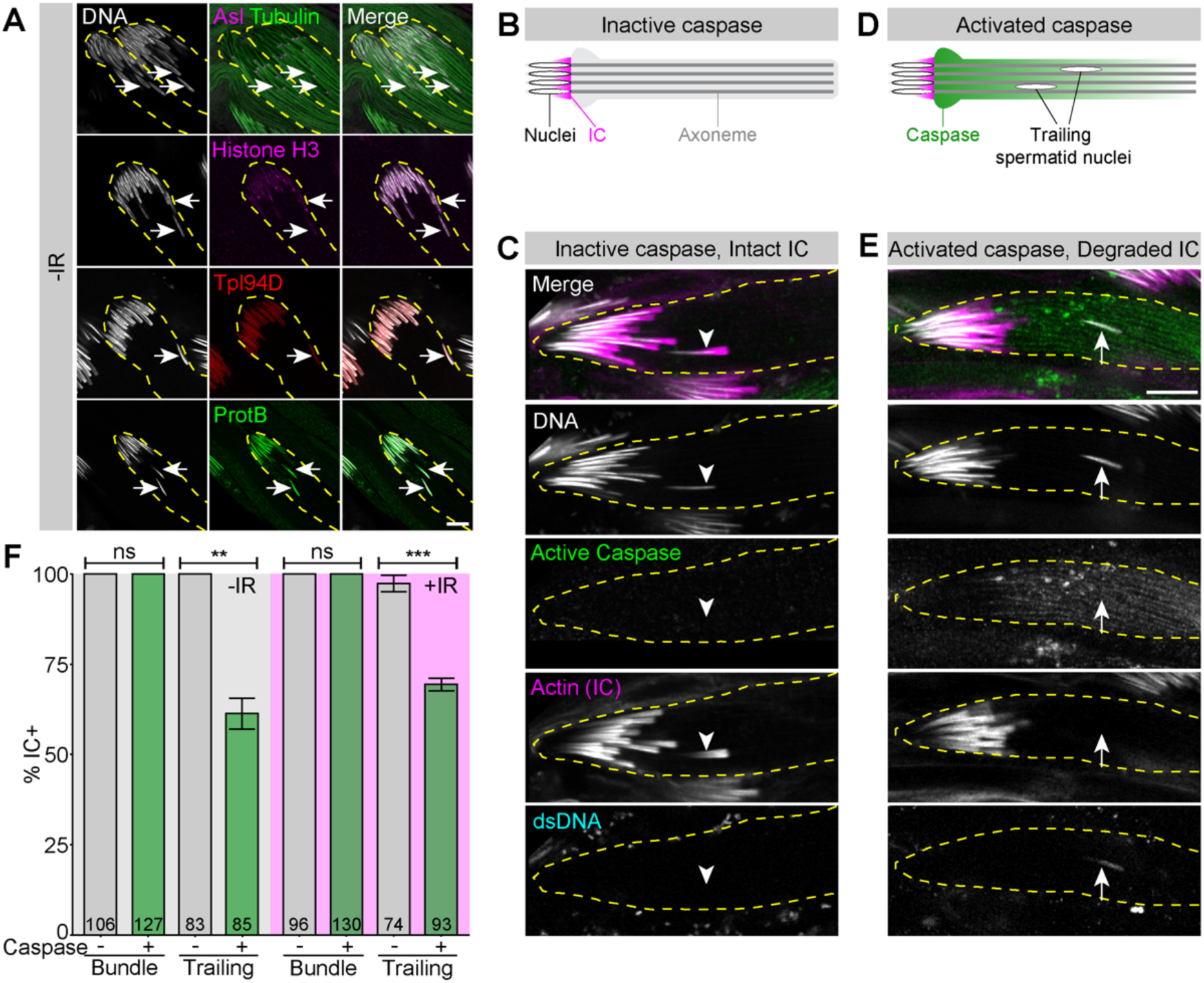
Trailing nuclei are eliminated by individualization. **A.** Representative images of spermatid nuclei stained for various markers of spermatid morphology (Asl in magenta, Tubulin in green), and histone-to-protamine exchange (Histone H3 in pink, Tpl94D in red, Protamine B in green). **B, D.** Schematic of individualization before (B) and upon (D) caspase activation. **C, E.** Representative image of spermatid cyst before (C) and after caspase activation (E) with trailing nuclei marked with an arrowhead (C) and arrow (E). Immunofluorescence of nuclei (DAPI, white), activated caspase (green), ICs (phalloidin, magenta), and dsDNA (cyan). Antibodies cannot penetrate tightly compacted needle stage nuclei, thus α-dsDNA serves as a marker for nuclear compaction. **F.** Quantification of IC+ nuclei in caspase-negative (gray bars) and caspase-positive (green bars) cysts without (gray background) and with irradiation (magenta background). The means with 95% CIs are plotted. N denotes number of spermatid bundles or individual trailing nuclei. For all images, yellow dashed lines outline the cyst in each image. Scale = 10 µm.

Importantly, we find that irradiation does not impact whether or not the IC is removed from trailing nuclei (Fig. 3F). However, irradiation increases trailing nuclei (Fig. 1C-E, 2), leading to an overall increase in the elimination of spermatid nuclei upon irradiation. Taken together, these results reveal that spermatid nuclei are eliminated upon irradiation via induced trailing and subsequent failure during individualization. We propose that nuclear trailing is the mechanism to eliminate subpar haploid spermatids more generally.

### Early spermatids exhibit elevated γH2Av upon irradiation

The above results suggest that trailing of nuclei serves as a mechanism to eliminate nuclei with damaged genomes, thereby ensuring that only high-quality genomes are passed to the offspring. These results also suggest the presence of a mechanism within haploid nuclei that senses DNA damage after the genomic insult and elicits the trailing response. In somatic diploid cells, the histone variant H2Av in *Drosophila* (H2Ax in mammals) is the primary mediator of DNA damage response through phosphorylation at S137 (denoted as γH2Av / γH2Ax) (Madigan et al., 2002; Rogakou et al., 1998; Van Daal and Elgin, 1992). It is well established that H2Av / H2Ax are phosphorylated by ATM, ATR, and DNA-PK kinases at the sites of DNA double-strand breaks, triggering downstream responses including repair and cell death (Ward and Chen, 2001; Burma et al., 2001; Ciccia and Elledge, 2010).

Given its universal role, we tested whether the spermatid response to DNA damage may involve γH2Av, although the spectrum of responses downstream of γH2Av / γH2Ax would be different in haploid spermatids. H2Av expression was observed in the germ cell nuclei throughout spermatogenesis, until it became undetectable around canoe stage during the histone-to-protamine transition, which removes the majority of histones from the paternal genome (Fig. 4A) (Rathke et al., 2014, 2007). Without irradiation, spermatid nuclei of round and leaf stages, where chromatin is still packaged by histones, showed very little γH2Av signal, but γH2Av became clearly detectable upon irradiation (Fig. 4B-E), suggesting that spermatids of these stages are responding to irradiation-induced DNA damage. A robust γH2Av signal was observed during the late canoe stage even without irradiation, likely reflecting known extensive DNA double-stranded breaks that are reported to facilitate the histone-to-protamine transition (Fig. S2A-C) (Marcon and Boissonneault, 2004; Rathke et al., 2007; Akematsu et al., 2017; Grégoire et al., 2018; Laberge and Boissonneault, 2005). Because of this strong γH2Av signal without irradiation, we could not assess the irradiation-specific response of the canoe-stage spermatids. However, γH2Av signal of early canoe spermatids (prior to a strong γH2Av signal) was similar between irradiated and non-irradiated testes (Fig. S2A, B), implying that canoe spermatids may not respond to DNA damage via γH2Av as much as earlier spermatids. These results suggest that haploid spermatids, in particular during the round and leaf stages, may respond to DNA damage via γH2Av-mediated signaling.

**Figure 4.**
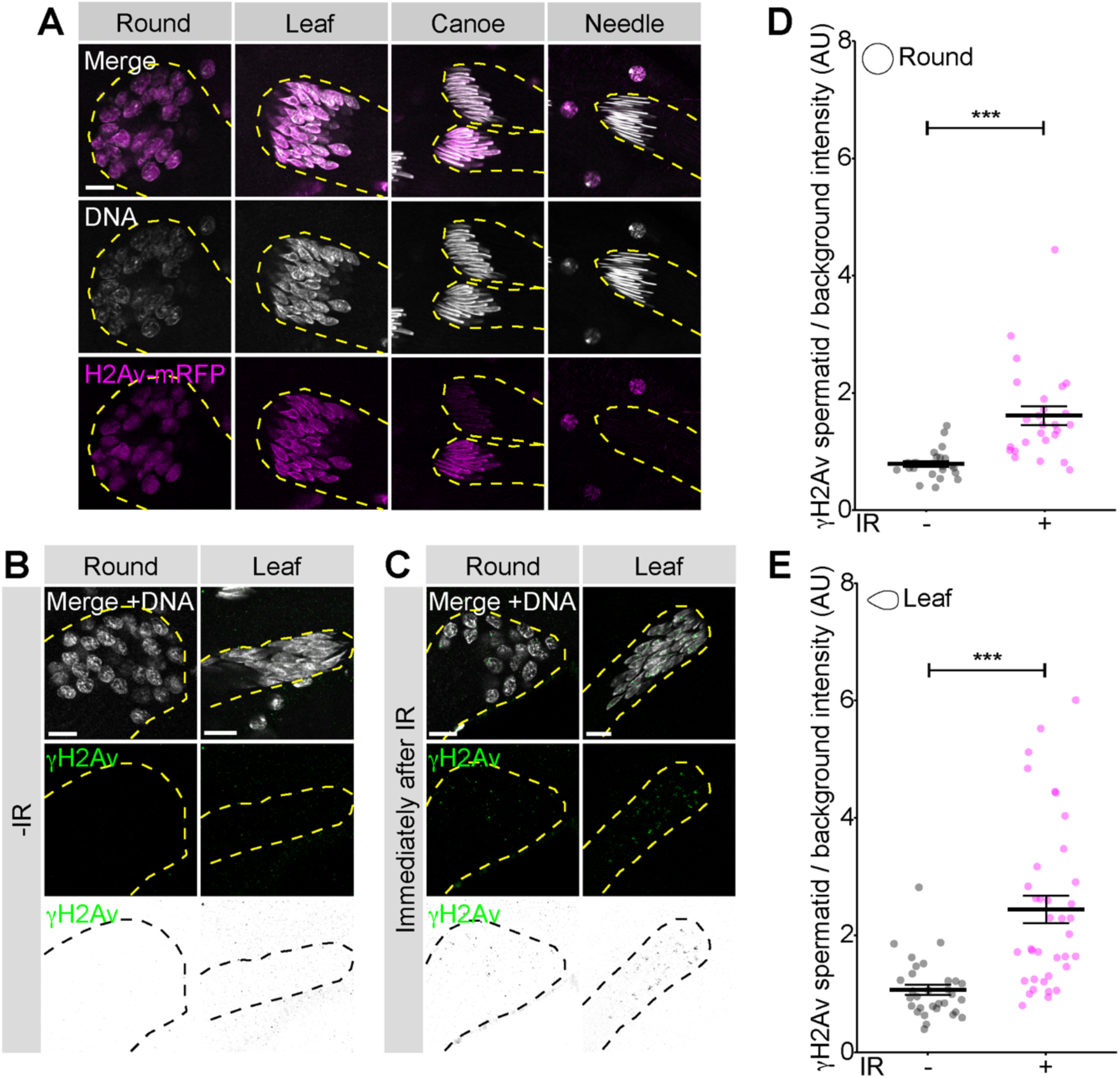
Spermatids contain histone variant H2Av and signal DNA damage through γH2Av. **A.** Representative images of H2Av-mRFP (magenta) localization to each spermatid nuclei stage (DAPI, white). **B-C.** Representative images of γH2Av (green, middle panels, and inverted LUT, bottom panels) localization on round and leaf spermatid nuclei (DAPI, white) without (B) and immediately after (C) irradiation. **D-E.** Quantification of γH2Av intensity in round (D) and leaf (E) nuclei without (gray) and immediately after IR (magenta), normalized to the background intensity. The means with 95% CIs are plotted. For all images, yellow dashed lines outline the cyst in each image. Scale = 10 µm.

### Histone H2Av is required for sperm nuclei trailing upon irradiation

To understand the role of histone H2Av in the DNA damage response of haploid spermatids, we used RNAi-mediated knockdown of H2Av in the male germline using the *bam-GAL4* driver (*bam-GAL4*>*UAS-RNAi*) to specifically deplete H2Av in the late spermatogonia to spermatocyte stage (Chen and McKearin, 2003). Knockdown efficiency was confirmed by the loss of H2Av-mRFP signal from a tagged transgenic strain (*H2Av-mRFP* (Schuh et al., 2007)) (Fig. S3A-C). We used two independent RNAi lines (*H2Av^HMS00162^* or *H2Av^HMS02773^*). Both RNAi lines depleted H2Av efficiently, with *H2Av^HMS00162^* displaying a more robust knockdown (Fig. S3A-C), thus we used *H2Av^HMS00162^* for further experiments. *H2Av^RNAi^*testes showed no γH2Av signal even after irradiation (Fig. S4A-D), further confirming efficient depletion of the γH2Av response. Unexpectedly, depletion of H2Av from late spermatogonia and onward caused no major visible defects on germ cell development (Fig. S3A, B), and *H2Av^RNAi^*males are indeed fertile with the seminal vesicles containing abundant mature sperm (Fig. S3D). These results demonstrate that H2Av is dispensable for germ cell development from spermatocytes through spermiogenesis stages.

Strikingly, however, *H2Av^RNAi^* testes exhibited a marked decrease in spermatid trailing upon irradiation (Fig. 5A, B). Counting basal bodies in the nuclear bundle (stained with Asl, as in Fig. 2) confirmed that the irradiation-induced loss of nuclei did not occur upon RNAi-mediated knockdown of H2Av (Fig. 5C, D). Together, these data suggest that γH2Av-mediated signaling is responsible for nuclear trailing upon irradiation. In the absence of γH2Av signaling, DNA-damaged spermatids remain in the main bundle instead of trailing.

**Figure 5.**
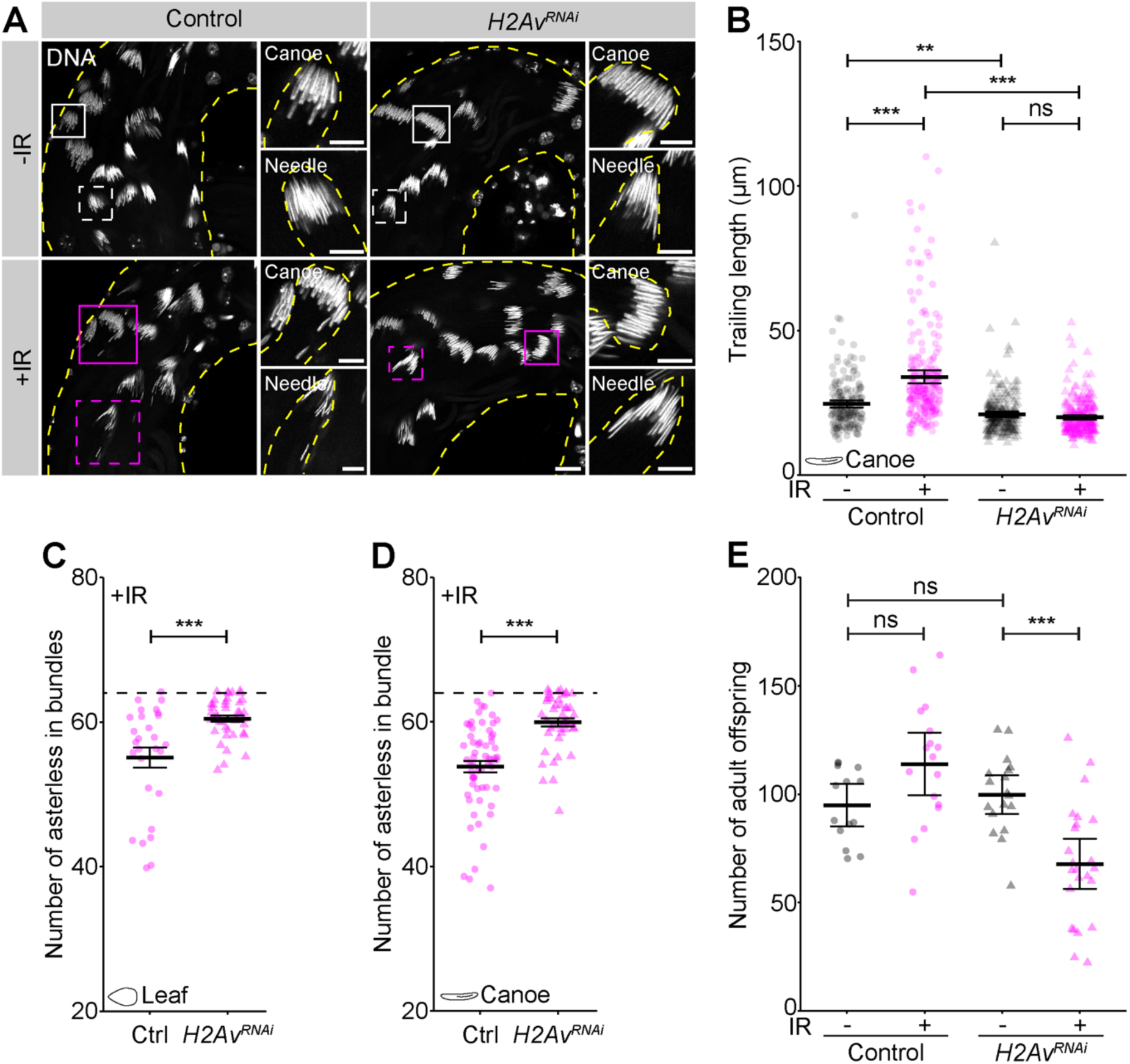
H2Av is required for spermatid trailing in response to DNA damage. **A.** Representative images of spermatid nuclei stained with DAPI (white) in control vs. *H2Av^RNAi^* testes without (top panels) and with irradiation (bottom panels) at 1 dpIR, scale = 20 µm. Inset images show single canoe (top, solid box) and needle (bottom, dashed box) cysts, scale = 10 µm. Yellow dashed lines outline the testis or cyst outlines in each image. **B-E.** Quantification of spermatid cyst trailing length (B) and asterless count per cyst (C, D) at 1 dpIR, and progeny counts (E) across non-irradiated (gray) and irradiated (magenta) flies from control (circles) and *H2Av^RNAi^*(triangles) genotypes. In C-D, the dotted line marks 64, the theoretical maximum number of nuclei per cyst. The means with 95% CIs are plotted.

Because the nuclei in the bundle are protected from elimination, we hypothesized that damaged nuclei may be allowed to proceed with individualization and become sperm upon depletion of *H2Av*, contributing to fertilization. Intriguingly, we found that *H2Av^RNAi^* males exhibited reduced fertility upon irradiation, although *H2Av^RNAi^*alone did not affect fertility in the absence of irradiation (Fig. 5E). These results indicate that embryos fertilized by defective, DNA-damaged sperm fail to develop successfully. In support of this idea, we found a slight increase in the number of seminal vesicles that contained dsDNA-positive sperm nuclei after irradiation upon *H2Av* depletion, indicating that sperm with damaged DNA were indeed successfully individualized and became mature sperm (Fig. S5A, B). Together, these results suggest that H2Av is required for irradiation-induced spermatid trailing, preventing defective nuclei from becoming mature sperm, and thereby ensuring the quality of gametes.

### Phosphorylation of H2Av mediates haploid genome quality control

These results revealed the importance of H2Av in the DNA damage response of haploid spermatids. Because we observed that round and leaf spermatids induce phosphorylation of H2Av (i.e., γH2Av, phosphorylation at S137 (Rogakou et al., 1998)) (Fig. 4B-E), we next examined whether S137 plays a role in spermatids’ response to irradiation.

We generated transgenes that express non-phosphorylatable (*H2Av^S137A^*), phosphomimetic (*H2Av^S137E^*), and wild type (*H2Av^WT^*, control) H2Av under the *β2-tubulin* promoter, which drives expression in spermatocytes onward (Michiels et al., 1989). All constructs were successfully expressed in the testis, without any major effect on spermatogenesis. Strikingly, however, the expression of the *H2Av^S137A^* non-phosphorylatable mutant in the late germ cells led to an almost complete lack of trailing upon irradiation, while flies expressing *H2Av^WT^* transgene exhibited normal trailing (Fig. 6A-C). These results suggest that S137 phosphorylation is essential to induce trailing upon irradiation.

**Figure 6.**
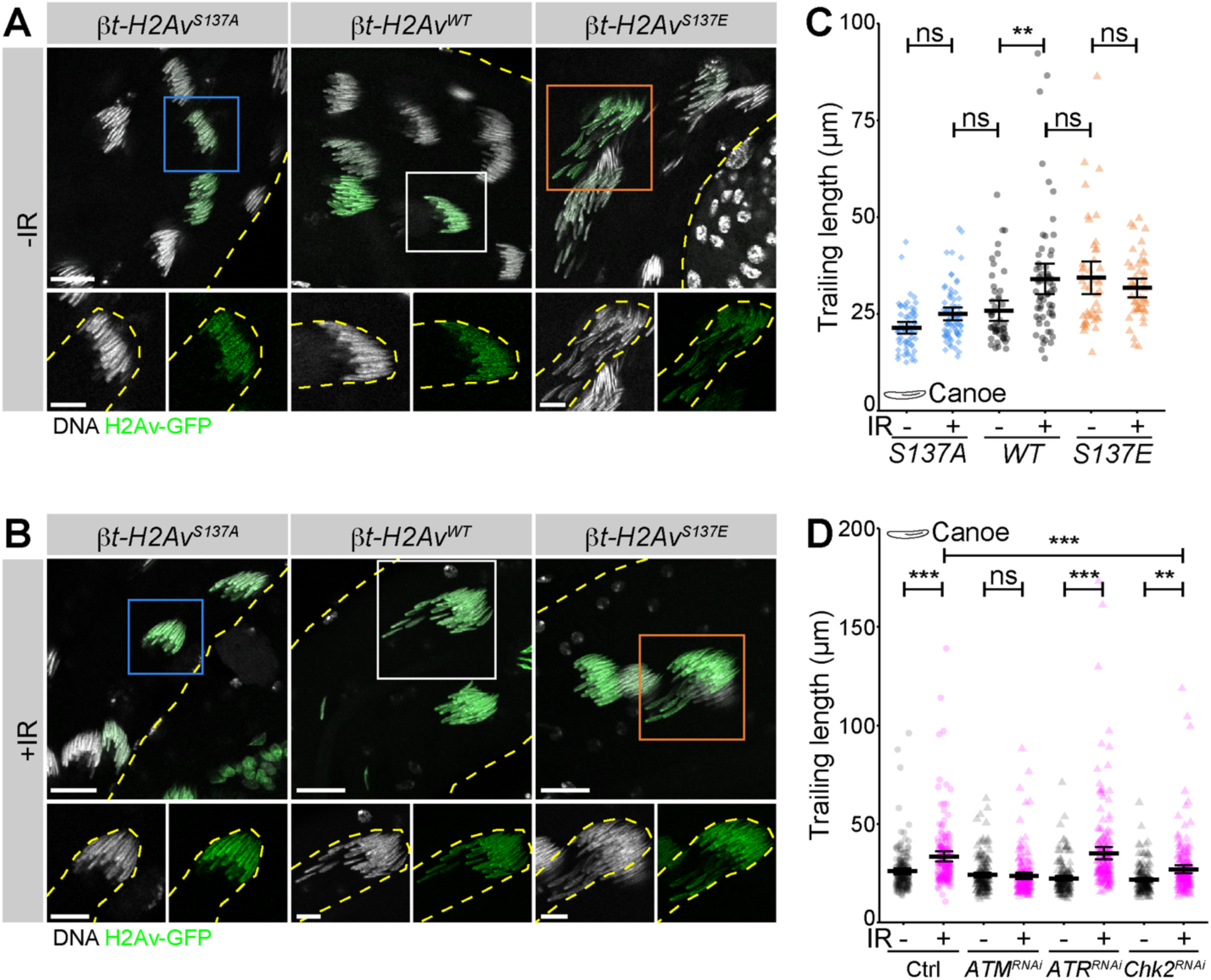
H2Av S137 is required for spermatid trailing in response to DNA damage. **A-B.** Representative images of spermatid nuclei in *H2Av^WT^* vs. *H2Av^S137^* phospho-variants without (A) and with (B) irradiation at 1 dpIR, scale = 20 µm. Inset images show single canoe cysts, scale = 10 µm. Yellow dashed lines outline the testis or cyst in each image. DAPI is shown in white, βt-H2Av-GFP in green. **C.** Quantification of spermatid cyst trailing length without and with irradiation across *H2Av^S137^* phospho-variant genotypes (S137A in blue diamonds, WT in gray circles, S137E in orange triangles). The means with 95% CIs are plotted. **D.** Quantification of spermatid cyst trailing length without (gray) and with irradiation (magenta) across control (circles) and RNAi lines (triangles). The means with 95% CIs are plotted.

Conversely, expression of the *H2Av^S137E^* phosphomimetic transgene led to a dramatic increase in trailing, even in the absence of irradiation. Moreover, testes expressing *H2Av^S137E^* did not further increase trailing upon irradiation (Fig. 6A-C), suggesting that irradiation-induced trailing may solely rely on *H2Av^S137E^*. Collectively, these data suggest that the *H2Av* S137 phosphorylation site controls spermatid nuclei trailing in response to DNA damage, and represents a non-canonical function for H2Av and γH2Av signaling.

H2Av phosphorylation is normally mediated by conserved DNA damage response kinases, ATM and ATR, both of which can directly phosphorylate H2Av. ATM is the primary kinase activated in response to double-stranded breaks, such as those caused by ionizing radiation, while ATR functions predominantly during replicative stress (Joyce et al., 2011; Ward and Chen, 2001; Burma et al., 2001). To understand their potential contribution to H2Av phosphorylation, we examined irradiation-induced nuclear trailing when these kinases were depleted using previously validated RNAi lines (see Methods). We found that *ATM^RNAi^* testes, but not *ATR^RNAi^* testes, showed a lack of trailing after irradiation. This suggests that H2Av is primarily phosphorylated by ATM to induce γH2Av-mediated trailing (Fig. 6D). ATM is known to phosphorylate effector kinase Chk2 to recruit downstream DNA repair components to the break site (Matsuoka et al., 1998, 2000; Zannini et al., 2014). *Chk2^RNAi^* testes showed a slight reduction in irradiation-induced trailing (Fig. 6D), indicating that spermatid trailing is primarily mediated by the ATM-H2Av signaling axis with Chk2 contributing partially to this response.

Together, these results suggest that haploid spermatids use a canonical DNA damage signaling pathway (ATM and γH2Av) to sense and signal DNA damage. However, interestingly, H2Av phosphorylation downstream of canonical DNA damage sensing leads to a haploid spermatid-specific response, nuclear trailing and spermatid elimination, ensuring genome quality of the spermatids that will participate in fertilization. It awaits future investigations how γH2Av is mechanistically linked to spermatid trailing.

## DISCUSSION

Despite being dispensable for the survival of the organism, gamete genome integrity is fundamental to allow reproductive success. Our study uncovers a DNA damage response that acts specifically in the vulnerable haploid stages of spermiogenesis, a uniquely challenging context for genome integrity.

The elevated sensitivity of germ cells to DNA damage is well known. Prior studies have demonstrated how diploid germ cells trigger a unique cell death program in response to irradiation-induced DNA damage rather than attempting DNA repair. Mitotically-dividing germ cells (spermatogonia) are extremely sensitive to DNA damage, where interconnected germ cells within a cyst communicate DNA damage and enhance cell death across the entire cyst. This response is thought to provide heightened stringency in germline genome quality control, though it commonly results in infertility after radiation therapy and chemotherapy (Lu and Yamashita, 2017; Edwards and Sirlin, 1958; Meistrich, 2013; Oakberg, 1955).

Haploid-stage (i.e., post-meiotic) germ cells face unique challenges in maintaining genome integrity, and the strategies employed by somatic cells or diploid-stage germ cells are unlikely to be effective. First, post-meiotic germ cells lack homologous chromosomes and sister chromatids that serve as templates for accurate DNA repair. Second, the reduced transcriptional capacity during post-meiotic spermiogenesis will limit the ability to respond to DNA damage. It has remained unknown if and how post-meiotic cells monitor genome quality and prevent subpar genomes from contributing to the next generation. This is the first study to describe the mechanism employed by haploid-stage germ cells to ensure the genome quality of gametes. We showed that irradiation induced ‘trailing’ of developing spermatids, leading to their eventual elimination. This suggests that spermatids employ a ‘sense damage and toss’ strategy, rather than attempting repair, a strategy especially suited for cells that lack a template for accurate DNA repair. We further demonstrate that γH2Av is the spermatids’ DNA damage sensor, which unexpectedly leads to a non-canonical response of ‘nuclear trailing’ and thus spermatid elimination.

The underlying mechanism that transmits γH2Av signal to lead to trailing remains unknown and awaits future investigation. It likely involves cytoskeletal mechanisms, such that nuclei become unable to remain in the main spermatid bundle. We note that trailing spermatids appear to have an intact basal body and attachment to the microtubule axoneme (Fig. 3A), indicating that detachment of the nuclei from the microtubule is not the underlying mechanism for the trailing, although other defects, such as lack of axonemal growth, remain possible. There are a few precedents that connect DNA damage to cytoskeletal mechanisms. For example, DNA damage can induce actin filaments within the nucleus in HeLa cells and oocytes, and this nuclear actin assembly promotes efficient DSB clearance (Belin et al., 2015; Sun et al., 2017). Additionally, centrosomes have been coupled to DNA damage-induced cell cycle defects and subsequent loss of the damaged cells. Upon irradiation or Rad51 deficiency, DT40 cells undergo centrosome amplification in a largely ATM-dependent manner during a prolonged G2 phase to eliminate cells with substantial DNA damage (Dodson et al., 2004). In *Drosophila* embryos, double-strand breaks trigger Chk2-dependent centrosome inactivation, leading to mitotic failure and nuclear elimination (Takada et al., 2003). DNA damage proteins, including ATM, ATR, and Chk2, localize to centrosomes as well (Takada et al., 2003; Mullee and Morrison, 2016). It is possible that similar mechanisms may also link γH2Av DNA damage signaling and cytoskeletal changes in spermatid trailing.

Altogether, we identify that H2Av mediates spermatid nuclear trailing and elimination in response to DNA damage. We show that trailing nuclei are eliminated by failed individualization, and this mechanism could prevent defective spermatids, DNA-damaged or otherwise, from being inherited by the next generation, ultimately protecting reproduction.

## ACKNOWLEDGEMENTS

We thank the Yamashita lab members, past and present, for discussions and expertise. We are grateful to Flybase and the Bloomington *Drosophila* Stock Center for reagents and resources. MK thanks the HHMI Hanna Gray Fellows, Leading Edge Fellows, and Mark J. Khoury for community support and expertise. This research was supported by an HHMI Hanna H. Gray fellowship (to MK), and the Howard Hughes Medical Institute and Gordon and Betty Moore Foundation (to YMY).

## AUTHOR CONTRIBUTIONS

MK and YMY conceived the study. MK designed, performed, and analyzed all experiments. YMY supervised the study. MK and YMY wrote the manuscript.

## METHODS

### Fly husbandry and strains

All fly stocks were raised on standard Bloomington medium without propionic acid at 25°C, and young flies (1-3-day-old adults) were used for all experiments. Flies used for wildtype experiments were the standard lab wildtype strain *yw*. For all other experiments, control flies were siblings from the same genetic cross, such as in Fig. S3. The following fly stocks were used: *H2Av-mRFP* (Bloomington *Drosophila* Stock Center [BDSC]: 23651), *bam-gal4* (BDSC: 80579), *H2Av^HMS00162^* (BDSC: 34844), *H2Av^HMS02773^* (BDSC: 44056), *ATM^GL00138^* (BDSC: 44417), *ATR^GL00284^* (BDSC: 35371), *Chk2^GL00020^* (NIG: GL00020), *Tpl94D-mRFP* (Kyoto *Drosophila* Stock Center, DGRC 109817), *ProtB-GFP* (BDSC: 58406). *ATM^GL00138^*, *ATR^GL00284^*, and *Chk2^GL00020^* RNAi lines were previously validated (Sopko et al., 2014; Cosolo et al., 2019; Ma et al., 2016). We used FlyBase to find information on gene sequences and stocks.

### Irradiation

Irradiation experiments were performed with a Cesium-137 source with a dose rate of approximately 54-58 rad per minute over the course of this study. Young males were collected on the first day after eclosing and allowed to mate with females for ∼2 days at 25°C prior to irradiation. Flies were dissected and processed for immunofluorescence (below) at various time points after irradiation.

### Immunofluorescence and imaging

Testes were dissected in 1x PBS, transferred to 4% formaldehyde in 1x PBS, and fixed for 30 minutes. Fixed testes were then washed twice in 1x PBST (1x PBS + 0.1% Triton X-100) for at least 1 h, and blocked for 1 h in 1x PBST + 3% BSA. Samples were incubated with primary antibodies in fresh 1x PBST + 3% BSA overnight at 4°C. Testes were washed thrice in 1x PBST for at least 1 h total, followed by incubation with secondary antibodies overnight at 4°C in 1x PBST + 3% BSA. Following 3 washes in 1x PBST, samples were mounted in Vectashield with DAPI (Vector Labs). Primary antibodies used: anti-Asl (1:5000, generously gifted from Tomer Avidor-Reiss), anti-ATP5a (1:1000, Abcam ab14748), anti-hts 1B1 (adducin, 1:20, DSHB), anti-Histone H3 (1:250, Abcam ab1791), anti-Cleaved caspase 3 (1:100, Cell Signaling Technology), anti-dsDNA (1:200, Abcam ab27156), anti-γH2Av (1:200, Rockland). Secondary antibodies used: Phalloidin-Alexa Fluor 568 and 647 (1:200; Thermo Fisher Scientific), FITC-tubulin (1:200; Thermo Fisher Scientific). Alexa Fluor-conjugated secondary antibodies were all used at a 1:200 dilution. Testes were imaged on an inverted Leica Stellaris 8 confocal microscope with a 40x/NA 1.3 or 63x/NA 1.4 immersion oil objective. Images were processed using FIJI/ImageJ software (below).

### Image Analysis

All analysis was done in FIJI/ImageJ.

Trailing length is defined as a segmented line starting from the head/acrosome-side of a spermatid bundle and following the curve of the cyst until the last visible trailing nucleus. Only cysts where trailing nuclei could be confidently traced were measured. Importantly, z-stacks were acquired to enable this nuclear tracing through longer cysts.

Basal bodies were marked and counted using an anti-Asl antibody. Z-stacks of individual bundles were acquired, and maximum projections were created to enable accurate basal body counting. Any spermatid bundles that overlapped with a neighboring bundle in any dimension were not included in this analysis. As we were interested in determining the number of nuclei that remain in the spermatid bundle, we used the position of nuclei to first define visibly trailing nuclei and exclude those basal bodies (Fig. 2B).

To measure γH2Av intensity, we identified a central z-plane within a spermatid bundle and created an ROI encompassing the entire bundle. This γH2Av measurement was normalized against the background intensity for the same-sized ROI.

Germline H2Av knockdown efficiency was measured in single spermatid nuclei, where the intensities for H2Av-mRFP were normalized against DAPI and compared against somatic nuclei using the same ROI.

### Fertility assay

For each genotype, 2 independent experiments consisting of 10 males each were assayed as follows: Individual 0-day old males were crossed with three 1-4-day old virgin *yw* females for 5 days at 25°C. On the fifth day, females were discarded, and males were transferred to new vials with three fresh 1-4-day old virgin *yw* females. All F1 progenies were counted from each vial to determine the number and sex ratio of the progeny. 0-day old males were irradiated as specified above and then immediately placed with virgin *yw* females. For each cross, the assay was discarded if the male or 2/3 females died during the course of the assay.

### Transgenic fly construction

The *β2-tubulin-H2Av-GFP* transgenic strains were generated via phiC31 site-directed integration into the *Drosophila melanogaster* genome. Constructs were synthesized by Vectorbuilder with the *β2*-tubulin promoter, and inserted into the attP40 integration site on the 2^nd^ chromosome by BestGene Inc.

## Supplementary Figures

**Figure S1.**
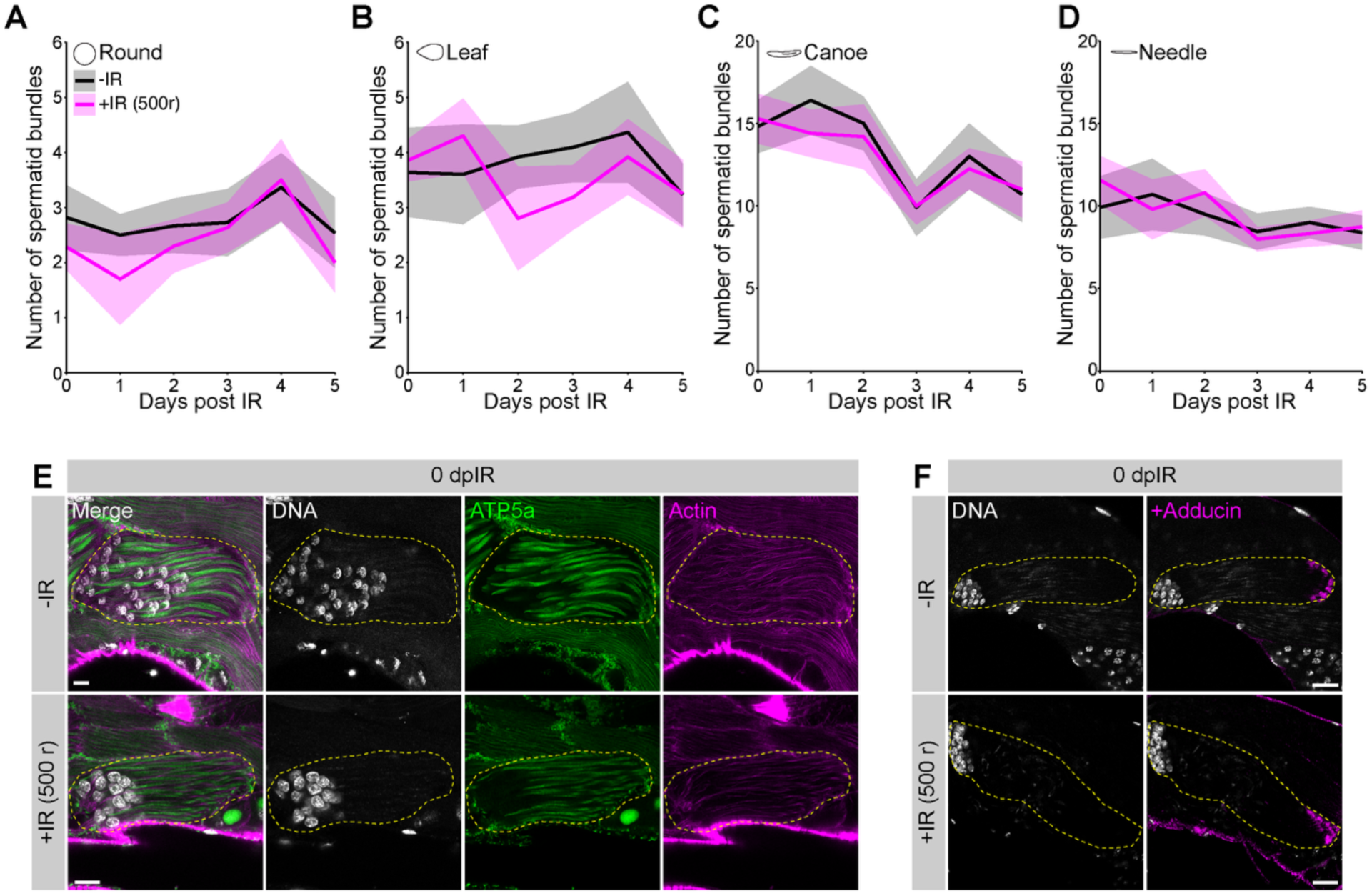
Irradiation does not lead to a spermiogenesis arrest or cyst disorganization. A-D. Quantification of the number of spermatid bundles present in non-irradiated (black) and irradiated (500 rad, magenta) flies across spermiogenesis nuclear stages. The means with 95% CIs are plotted from N > 10 testes per timepoint. **E, F.** Representative images of round spermatid nuclei cysts without (top) and immediately after irradiation (bottom), stained with DAPI (white), ATP5a (E, green), and actin (E, magenta) to show mitochondrial elongation, or adducin (F, magenta) to mark the distal end of the cyst. Yellow dashed lines outline the cyst in each image. Scale bars are 10 (E) and 20 µm (F).

**Figure S2.**
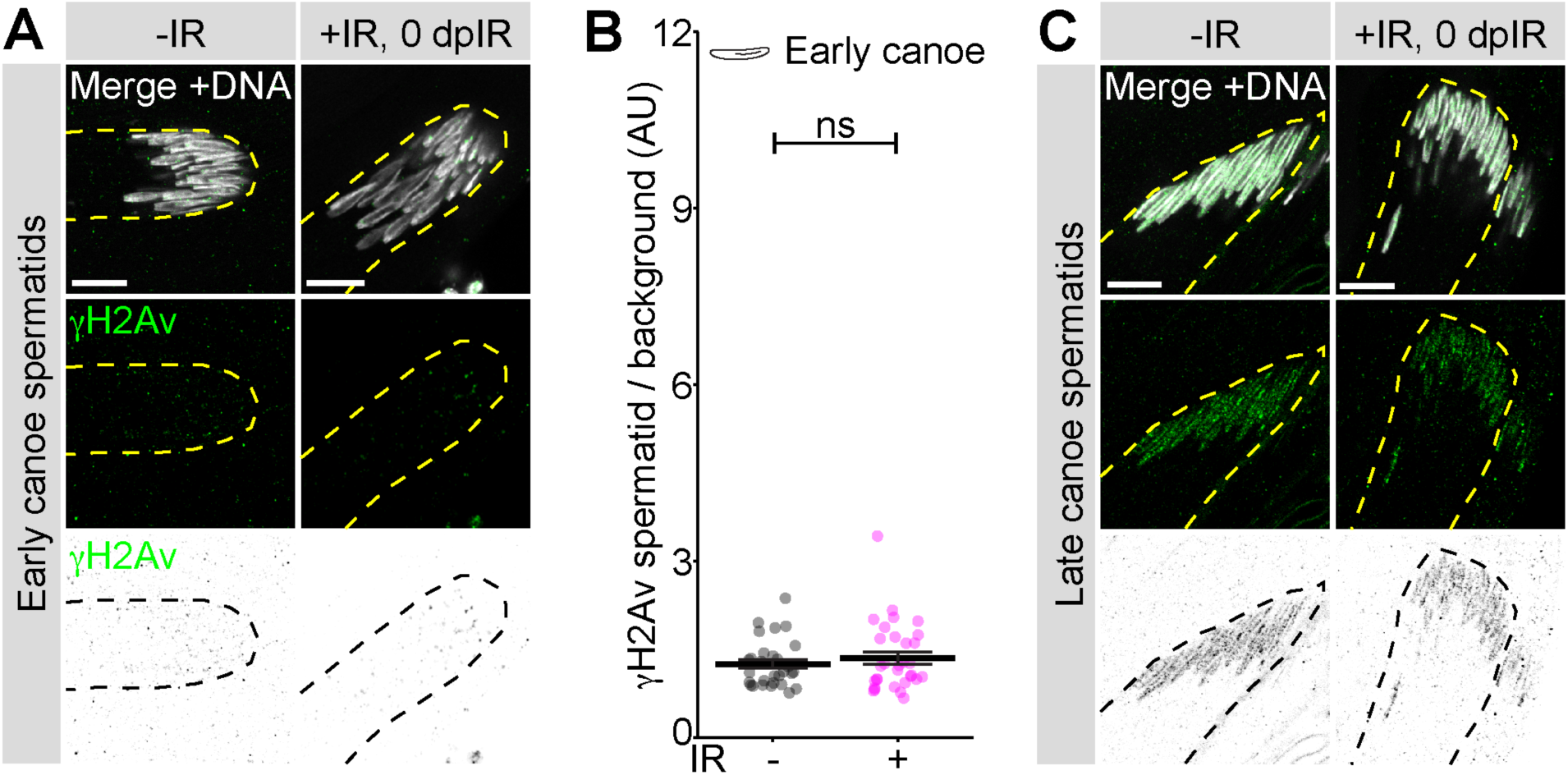
γH2Av localization in canoe stages. A, **C.** Representative images of γH2Av localization in early (A) and late (C) canoe spermatid stages without and immediately after irradiation. Needle nuclei are so tightly compacted that antibodies cannot penetrate and thus are not shown here. DAPI is shown in white, α-γH2Av is shown in green (middle panels) and as an inverted LUT (bottom panels). Yellow or black dashed lines outline the cyst in each image, scale = 10 µm. **B.** Quantification of γH2Av intensity in early canoe nuclei without (gray) and immediately after IR (magenta), normalized to the background intensity. The means with 95% CIs are plotted.

**Figure S3.**
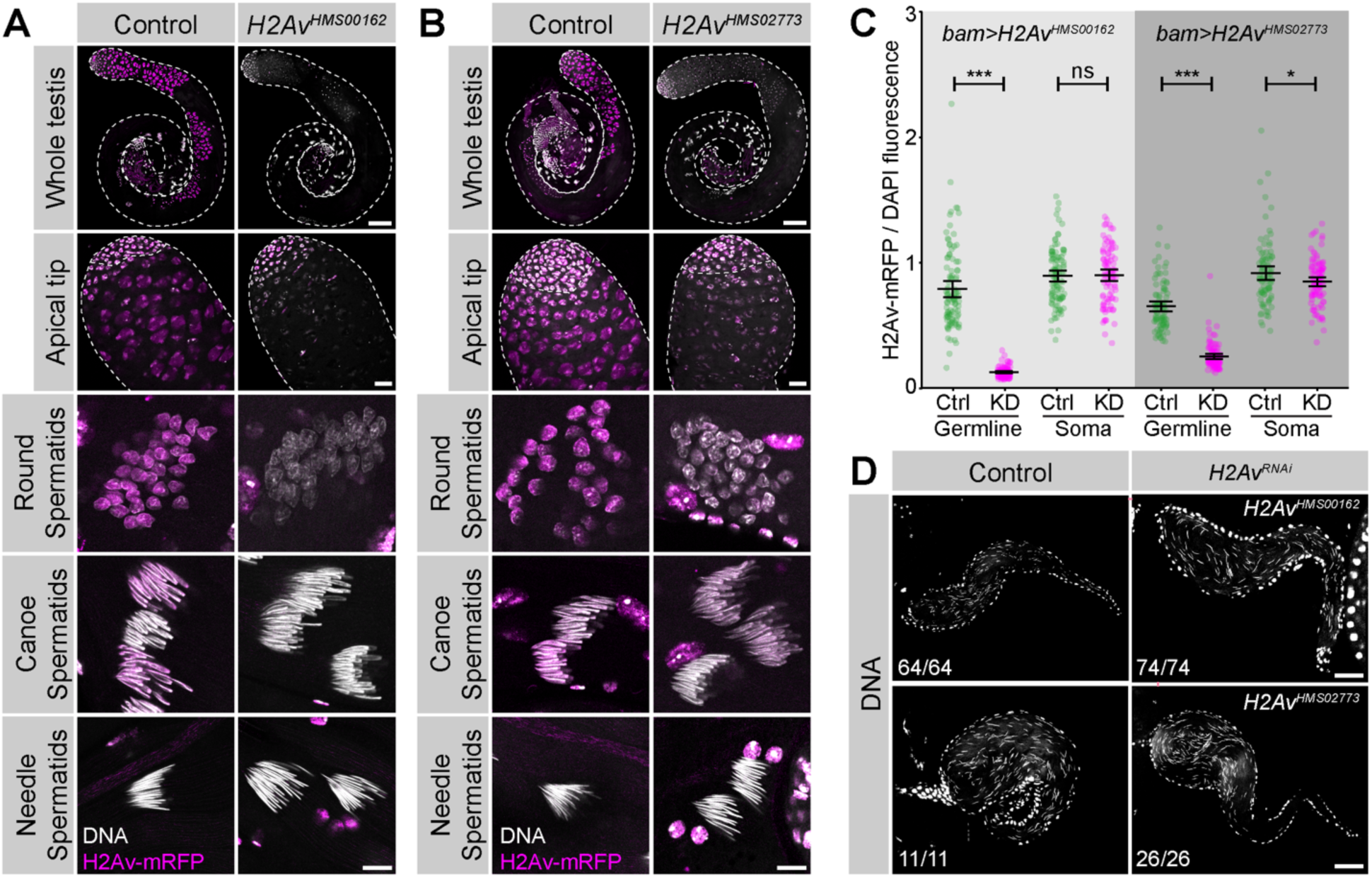
Two independent *H2Av^RNAi^* lines successfully knockdown H2Av. A-B. Representative images of control (left) and *H2Av^RNAi^* (right) testes with a fluorescent H2Av-mRFP transgene (magenta) and stained with DAPI (white). Note that knockdown was driven by *bam-gal4*, which begins from late spermatogonia (apical tip panels). White dashed lines outline the testis, scale = 100 µm (whole testis), 20 µm (apical tip), and 10 µm (spermatids). **C.** Quantification of testis-specific knockdown efficiency from 2 RNAi lines as measured by the ratio of H2Av-mRFP / DAPI in both germline and soma from sibling control (green) and RNAi (magenta) testes. The means with 95% CIs are plotted. **D.** Representative images of seminal vesicles from control (left) and RNAi (right) testes, stained with DAPI (white). Scale = 50 µm.

**Figure S4.**
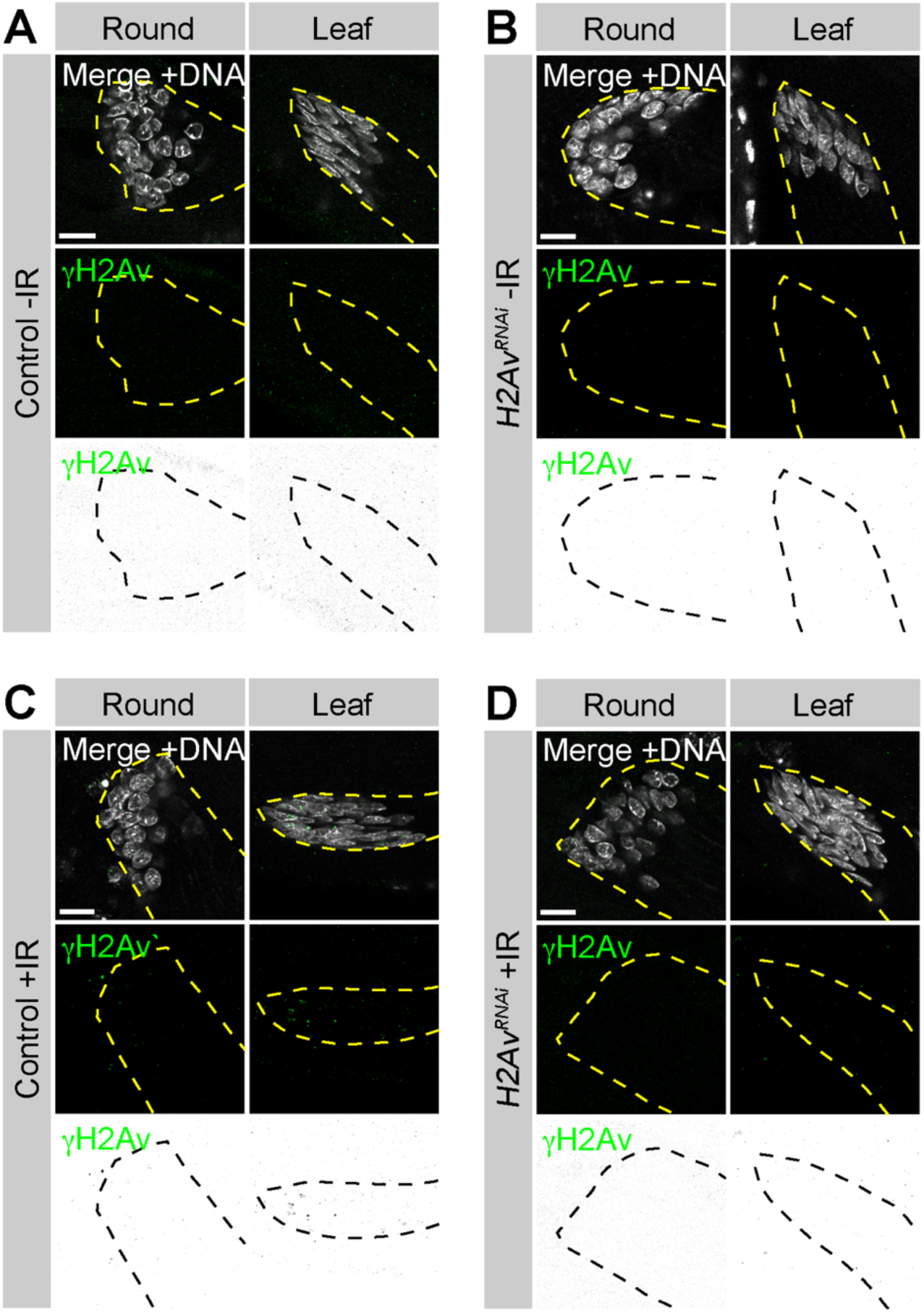
Knockdown of H2Av prevents spermatids from signaling with γH2Av after DNA damage. A-D. Representative images of control (A, C) and *H2Av^RNAi^* (B, D) spermatids in round and leaf stages without (A, B) and immediately after (C, D) irradiation. DAPI is shown in white, α-γH2Av is shown in green (middle panels) and as an inverted LUT (bottom panels). Yellow or black dashed lines outline the cyst in each image, scale = 10 µm.

**Figure S5.**
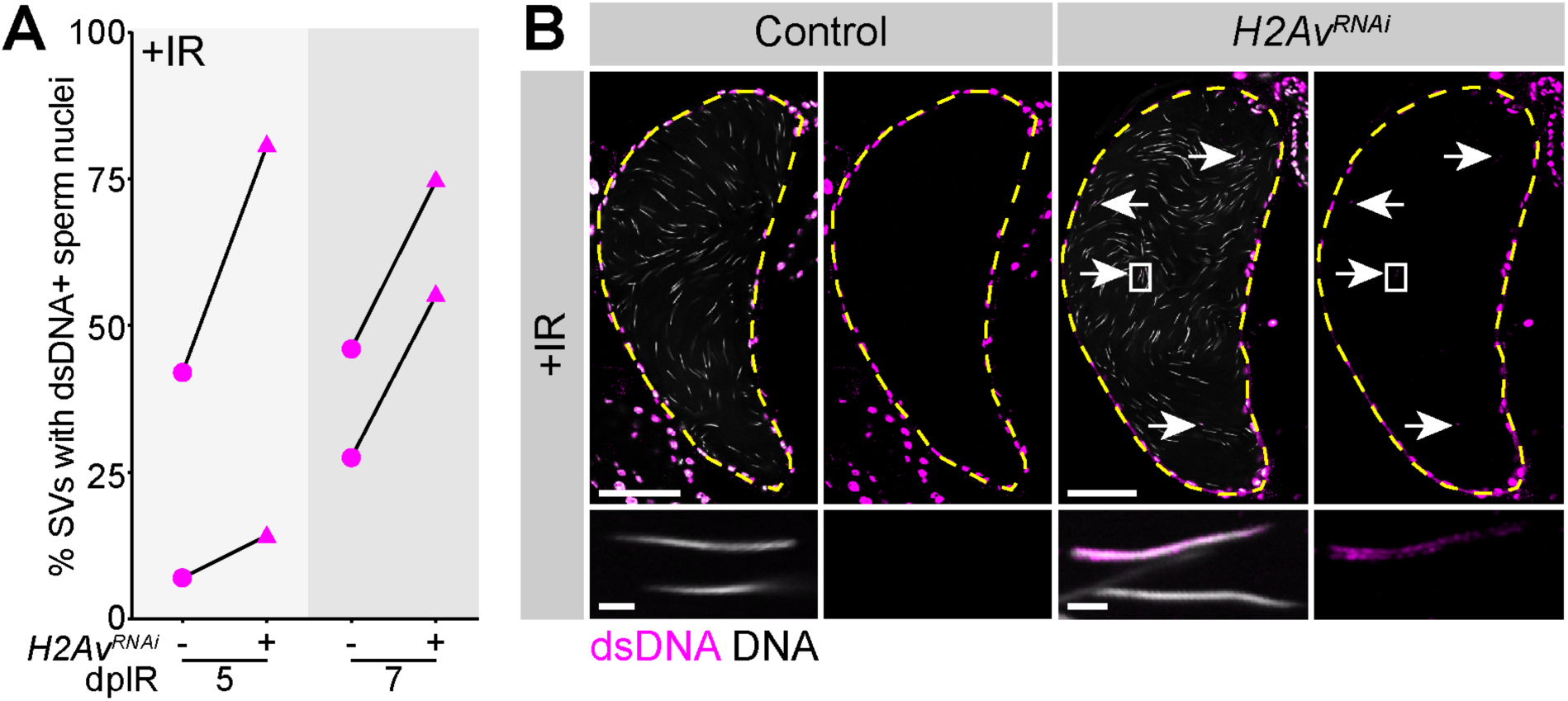
*H2Av^RNAi^* seminal vesicles show dsDNA+ sperm nuclei after irradiation. **A.** Quantification of control (circles) and *H2Av^RNAi^* (triangles) seminal vesicles with dsDNA+ sperm nuclei after irradiation. Black lines connect data points from the same experiment, with 2 replicates shown. **B.** Representative images of control and *H2Av^RNAi^*seminal vesicles after irradiation, stained with DAPI (white) and dsDNA (magenta). Arrows indicate dsDNA+ sperm nuclei, with one box for the inset image. Yellow dashed lines outline the seminal vesicle in each image, scale bar = 50 µm (top) and 2 µm (bottom, inset).

## REFERENCES

Akematsu, T., Y. Fukuda, J. Garg, J.S. Fillingham, R.E. Pearlman, and J. Loidl. 2017. Post-meiotic DNA double-strand breaks occur in Tetrahymena, and require Topoisomerase II and Spo11. Elife. 6:1–26. doi:10.7554/eLife.26176.

Arama, E., J. Agapite, and H. Steller. 2003. Caspase activity and a specific cytochrome C are required for sperm differentiation in Drosophila. Dev. Cell. 4:687–697. doi:10.1016/S1534-5807(03)00120-5.

Arama, E., M. Bader, M. Srivastava, A. Bergmann, and H. Steller. 2006. The two Drosophila cytochrome C proteins can function in both respiration and caspase activation. EMBO J. 25:232–243. doi:10.1038/sj.emboj.7600920.

Barreau, C., E. Benson, E. Gudmannsdottir, F. Newton, and H. White-Cooper. 2008. Post-meiotic transcription in Drosophila testes. Development. 135:1897–1902. doi:10.1242/dev.021949.

Belin, B.J., T. Lee, and R.D. Mullins. 2015. DNA damage induces nuclear actin filament assembly by formin-2 and spire-1/2 that promotes efficient DNA repair. Elife. 4:1–21. doi:10.7554/eLife.07735.

Buglak, D.B., K.H.M. Holmes, B.J. Galletta, and N.M. Rusan. 2024. The proximal centriole-like structure maintains nucleus-centriole architecture in sperm. J. Cell Sci. 137. doi:10.1242/jcs.262311.

Burma, S., B.P. Chen, M. Murphy, A. Kurimasa, and D.J. Chen. 2001. ATM phosphorylates histone H2AX in response to DNA double-strand breaks. J. Biol. Chem. 276:42462–42467. doi:10.1074/jbc.c100466200.

Chandley, A.C., and A.J. Bateman. 1962. Timing of Spermatogenesis in Drosophila melanogaster using Tritiated Thymidine. Nature. 193:299–300.

Chen, D., and D.M. McKearin. 2003. A discrete transcriptional silencer in the bam gene determines asymmetric division of the Drosophila germline stem cell. Development. 130:1159–1170. doi:10.1242/dev.00325.

Chen, M.Y., A. Tayyeb, and Y.F. Wang. 2021. shrub is required for spermatogenesis of Drosophila melanogaster. Arch. Insect Biochem. Physiol. 106:1–15. doi:10.1002/arch.21779.

Ciccia, A., and S.J. Elledge. 2010. The DNA Damage Response: Making It Safe to Play with Knives. Mol. Cell. 40:179–204. doi:10.1016/j.molcel.2010.09.019.

Cosolo, A., J. Jaiswal, G. Csordás, I. Grass, M. Uhlirova, and A.K. Classen. 2019. JNK-dependent cell cycle stalling in G2 promotes survival and senescence-like phenotypes in tissue stress. Elife. 8:1–27. doi:10.7554/eLife.41036.

Couderc, J.L., G. Richard, C. Vachias, and V. Mirouse. 2017. Drosophila LKB1 is required for the assembly of the polarized actin structure that allows spermatid individualization. PLoS One. 12:1–17. doi:10.1371/journal.pone.0182279.

Van Daal, A., and S.C.R. Elgin. 1992. Histone variant, H2AvD, is essential Drosophila melanogaster. Mol. Biol. Cell. 3:593–602. doi:10.1091/mbc.3.6.593.

Dodson, H., E. Bourke, L.J. Jeffers, P. Vagnarelli, E. Sonoda, S. Takeda, W.C. Earnshaw, A. Merdes, and C. Morrison. 2004. Centrosome amplification induced by DNA damage occurs during a prolonged G2 phase and involves ATM. EMBO J. 23:3864–3873. doi:10.1038/sj.emboj.7600393.

Dorogova, N. V., E.M. Akhmametyeva, S.A. Kopyl, N. V. Gubanova, O.S. Yudina, L. V. Omelyanchuk, and L.S. Chang. 2008. The role of Drosophila Merlin in spermatogenesis. BMC Cell Biol. 9:1–15. doi:10.1186/1471-2121-9-1.

Edwards, R.G., and J.L. Sirlin. 1958. The effect of 200 r of X-rays on the rate of spermatogenesis and spermiogenesis in the mouse. Exp. Cell Res. 15:522–528. doi:10.1016/0014-4827(58)90100-9.

Fabian, L., and J.A. Brill. 2012. Drosophila spermiogenesis. Spermatogenesis. 2:197–212. doi:10.4161/spmg.21798.

Fabrizio, J.J., G. Hime, S.K. Lemmon, and C. Bazinet. 1998. Genetic dissection of sperm individualization in Drosophila melanogaster. Development. 125:1833–1843. doi:10.1242/dev.125.10.1833.

Fatima, R. 2011. Drosophila dynein intermediate chain gene, Dic61b, is required for spermatogenesis. PLoS One. 6. doi:10.1371/journal.pone.0027822.

Graham, E.L., J. Fernandez, S. Gandhi, I. Choudhry, N. Kellam, and J.R. LaRocque. 2024. The impact of developmental stage, tissue type, and sex on DNA double-strand break repair in Drosophila melanogaster. PLoS Genet. 20:1–19. doi:10.1371/journal.pgen.1011250.

Grégoire, M.C., F. Leduc, M.H. Morin, T. Cavé, M. Arguin, M. Richter, P.É. Jacques, and G. Boissonneault. 2018. The DNA double-strand “breakome” of mouse spermatids. Cell. Mol. Life Sci. 75:2859–2872. doi:10.1007/s00018-018-2769-0.

Gunes, S., M. Al-Sadaan, and A. Agarwal. 2015. Spermatogenesis, DNA damage and DNA repair mechanisms in male infertility. Reprod. Biomed. Online. 31:309–319. doi:10.1016/j.rbmo.2015.06.010.

Heller, C.G., and Y. Clermont. 1963. Spermatogenesis in Man: An Estimate of Its Duration. Science (80-.). 140:184–186. doi:10.1126/science.140.3563.184.

Herbette, M., X. Wei, C.H. Chang, A.M. Larracuente, B. Loppin, and R. Dubruille. 2021. Distinct spermiogenic phenotypes underlie sperm elimination in the Segregation Distorter meiotic drive system. PLoS Genet. 17:1–26. doi:10.1371/journal.pgen.1009662.

Huh, J.R., S.Y. Vernooy, H. Yu, N. Yan, Y. Shi, M. Guo, and B.A. Hay. 2004. Multiple apoptotic caspase cascades are required in nonapoptotic roles for Drosophila spermatid individualization. PLoS Biol. 2. doi:10.1371/journal.pbio.0020015.

Joyce, E.F., M. Pedersen, S. Tiong, S.K. White-Brown, A. Paul, S.D. Campbell, and K.S. McKim. 2011. Drosophila ATM and ATR have distinct activities in the regulation of meiotic DNA damage and repair. J. Cell Biol. 195:359–367. doi:10.1083/jcb.201104121.

Khire, A., K.H. Jo, D. Kong, T. Akhshi, S. Blachon, A.R. Cekic, S. Hynek, A. Ha, J. Loncarek, V. Mennella, and T. Avidor-Reiss. 2016. Centriole Remodeling during Spermiogenesis in Drosophila. Curr. Biol. 26:3183–3189. doi:10.1016/j.cub.2016.07.006.

Kimura, S. 2013. The Nap family proteins, CG5017/Hanabi and Nap1, are essential for Drosophila spermiogenesis. FEBS Lett. 587:922–929. doi:10.1016/j.febslet.2013.02.019.

Kitaoka, M., and Y.M. Yamashita. 2024. Running the gauntlet: challenges to genome integrity in spermiogenesis. Nucleus. 15. doi:10.1080/19491034.2024.2339220.

Kracklauer, M.P., H.M. Wiora, W.J. Deery, X. Chen, B. Bolival, D. Romanowicz, R.A. Simonette, M.T. Fuller, J.A. Fischer, and K.M. Beckingham. 2010. The Drosophila SUN protein Spag4 cooperates with the coiled-coil protein Yuri Gagarin to maintain association of the basal body and spermatid nucleus. J. Cell Sci. 123:2763–2772. doi:10.1242/jcs.066589.

Laberge, R.-M., and G. Boissonneault. 2005. On the Nature and Origin of DNA Strand Breaks in Elongating Spermatids1. Biol. Reprod. 73:289–296. doi:10.1095/biolreprod.104.036939.

Lindsley, D.L., and K.T. Tokuyasu. 1980. Spermatogenesis. In Genetics and Biology of Drosophila (Ashburner, M. and Wright, T. R. F., eds.), Academic, New York. 225–294.

Lu, K.L., and Y.M. Yamashita. 2017. Germ cell connectivity enhances cell death in response to DNA damage in the Drosophila testis. Elife. 6:1–16. doi:10.7554/eLife.27960.

Ma, X., Y. Han, X. Song, T. Do, Z. Yang, J. Ni, and T. Xie. 2016. DNA damage-induced Lok/CHK2 activation compromises germline stem cell self-renewal and lineage differentiation. Dev. 143:4312–4323. doi:10.1242/dev.141069.

Madigan, J.P., H.L. Chotkowski, and R.L. Glaser. 2002. DNA double-strand break-induced phosphorylation of Drosophila histone variant H2Av helps prevent radiation-induced apoptosis. Nucleic Acids Res. 30:3698–3705. doi:10.1093/nar/gkf496.

Marcon, L., and G. Boissonneault. 2004. Transient DNA Strand Breaks during Mouse and Human Spermiogenesis: New Insights in Stage Specificity and Link to Chromatin Remodeling. Biol. Reprod. 70:910–918. doi:10.1095/biolreprod.103.022541.

Matsuoka, S., M. Huang, and S.J. Elledge. 1998. Linkage of ATM to cell cycle regulation by the Chk2 protein kinase. Science (80-.). 282:1893–1897. doi:10.1126/science.282.5395.1893.

Matsuoka, S., G. Rotman, A. Ogawa, Y. Shiloh, K. Tamai, and S.J. Elledge. 2000. Ataxia telangiectasia-mutated phosphorylates Chk2 in vivo and in vitro. Proc. Natl. Acad. Sci. U. S. A. 97:10389–10394. doi:10.1073/pnas.190030497.

Meistrich, M.L. 2013. Effects of chemotherapy and radiotherapy on spermatogenesis in humans. Fertil. Steril. 100:1180–1186. doi:10.1016/j.fertnstert.2013.08.010.

Michiels, F., A. Gasch, B. Kaltschmidt, and R. Renkawitz-pohl. 1989. A 14 bp promoter element directs the testis specificity of the Drosophila 32 tubulin gene. EMBO J. 8.

Mullee, L.I., and C.G. Morrison. 2016. Centrosomes in the DNA damage response—the hub outside the centre. Chromosom. Res. 24:35–51. doi:10.1007/s10577-015-9503-7.

Napoletano, F., B. Gibert, K. Yacobi-Sharon, S. Vincent, C. Favrot, P. Mehlen, V. Girard, M. Teil, G. Chatelain, L. Walter, E. Arama, and B. Mollereau. 2017. P53-Dependent Programmed Necrosis Controls Germ Cell Homeostasis During Spermatogenesis. PLoS Genet. 13:1–21. doi:10.1371/journal.pgen.1007024.

Nerusheva, O.O., N. V. Dorogova, N. V. Gubanova, O.S. Yudina, and L. V. Omelyanchuk. 2009. A GFP trap study uncovers the functions of Gilgamesh protein kinase in Drosophila melanogaster spermatogenesis. Cell Biol. Int. 33:586–593. doi:10.1016/j.cellbi.2009.02.009.

Noguchi, T., and K.G. Miller. 2003. A role of actin dynamics in individualization during spermatogenesis in Drosophila melanogaster. Development. 130:1805–1816. doi:10.1242/dev.00406.

Oakberg, E.F. 1955. Sensitivity and Time of Degeneration of Spermatogenic Cells Irradiated in Various Stages of Maturation in the Mouse. Radiat. Res. 2:369–391.

Rathke, C., W.M. Baarends, S. Awe, and R. Renkawitz-Pohl. 2014. Chromatin dynamics during spermiogenesis. Biochim. Biophys. Acta - Gene Regul. Mech. 1839:155–168. doi:10.1016/j.bbagrm.2013.08.004.

Rathke, C., W.M. Baarends, S. Jayaramaiah-Raja, M. Bartkuhn, R. Renkawitz, and R. Renkawitz-Pohl. 2007. Transition from a nucleosome-based to a protamine-based chromatin configuration during spermiogenesis in Drosophila. J. Cell Sci. 120:1689–1700. doi:10.1242/jcs.004663.

Rogakou, E.P., D.R. Pilch, A.H. Orr, V.S. Ivanova, and W.M. Bonner. 1998. DNA double-stranded breaks induce histone H2AX phosphorylation on serine 139. J. Biol. Chem. 273:5858–5868. doi:10.1074/jbc.273.10.5858.

Schuh, M., C.F. Lehner, and S. Heidmann. 2007. Incorporation of Drosophila CID/CENP-A and CENP-C into Centromeres during Early Embryonic Anaphase. Curr. Biol. 17:237–243. doi:10.1016/j.cub.2006.11.051.

Sopko, R., M. Foos, A. Vinayagam, B. Zhai, R. Binari, Y. Hu, S. Randklev, L.A. Perkins, S.P. Gygi, and N. Perrimon. 2014. Combining genetic perturbations and proteomics to examine kinase-phosphatase networks in drosophila embryos. Dev. Cell. 31:114–127. doi:10.1016/j.devcel.2014.07.027.

Steinhauer, J. 2015. Separating from the pack: Molecular mechanisms of Drosophila spermatid individualization. Spermatogenesis. 5:1–11. doi:10.1080/21565562.2015.1041345.

Sun, M.H., M. Yang, F.Y. Xie, W. Wang, L. Zhang, W. Shen, S. Yin, and J.Y. Ma. 2017. DNA double-strand breaks induce the nuclear actin filaments formation in cumulus-enclosed oocytes but not in denuded oocytes. PLoS One. 12:1–10. doi:10.1371/journal.pone.0170308.

Takada, S., A. Kelkar, and W.E. Theurkauf. 2003. Drosophila checkpoint kinase 2 couples centrosome function and spindle assembly to genomic integrity. Cell. 113:87–99. doi:10.1016/S0092-8674(03)00202-2.

Tokuyasu, K.T. 1972. Dynamics of spermiogenesis in Drosophila melanogaster. I. Individualization Process. J. Ultrasructure Res. 124:479–506. doi:10.1016/S0022-5320(75)80089-X.

Tokuyasu, K.T. 1974. Dynamics of spermiogenesis in Drosophila melanogaster. IV. Nuclear Transformation. J. Ultrastruct. Res. 48:284–303. doi:10.1016/S0022-5320(75)90013-1.

Tokuyasu, K.T. 1975. Dynamics of spermiogenesis in Drosophila melanogaster. VI. Significance of “onion” nebenkern formation. J. Ultrasructure Res. 53:93–112. doi:10.1016/S0022-5320(75)80089-X.

Ward, I.M., and J. Chen. 2001. Histone H2AX Is Phosphorylated in an ATR-dependent Manner in Response to Replicational Stress. J. Biol. Chem. 276:47759–47762. doi:10.1074/jbc.c100569200.

White-Cooper, H. 2010. Molecular mechanisms of gene regulation during Drosophila spermatogenesis. Reproduction. 139:11–21. doi:10.1530/REP-09-0083.

Yuan, X., H. Zheng, Y. Su, P. Guo, X. Zhang, Q. Zhao, W. Ge, C. Li, Y. Xi, and X. Yang. 2019. Drosophila Pif1A is essential for spermatogenesis and is the homolog of human CCDC157, a gene associated with idiopathic NOA. Cell Death Dis. 10. doi:10.1038/s41419-019-1398-3.

Zannini, L., D. Delia, and G. Buscemi. 2014. CHK2 kinase in the DNA damage response and beyond. J. Mol. Cell Biol. 6:442–457. doi:10.1093/jmcb/mju045.

